# A complex virome that includes two distinct emaraviruses is associated to virus-like symptoms in *Camellia japonica*

**DOI:** 10.1101/822254

**Authors:** C. Peracchio, M. Forgia, M. Chiapello, M. Vallino, M. Turina, M. Ciuffo

## Abstract

*Camellia japonica* plants manifesting a complex and variable spectrum of viral symptoms like chlorotic ringspots, necrotic rings, yellowing with necrotic rings, yellow mottle, leaves and petals deformations, flower color-breaking were studied since 1940 essentially through electron microscopic analyses; however, a strong correlation between symptoms and one or more well characterized viruses was never verified. In this work samples collected from symptomatic plants were analyzed by NGS technique and a complex virome composed by viruses members of the *Betaflexiviridae* and *Fimoviridae* families was identified. In particular, the genomic fragments typical of the emaravirus group were organized in the genomes of two new emaraviruses species, tentatively named Camellia japonica associated emaravirus 1 and 2. They are the first emaraviruses described in camellia plants and were always found solely in symptomatic plants. On the contrary, in both symptomatic and asymptomatic plants, we detected five betaflexiviruses isolates that, based on aa identitiy comparisons, can be classified in two new putative species called Camellia japonica associated betaflexivirus 1 and Camellia japonica associated betaflexivirus 2. Together with other recently identified betaflexiviruses associated to *Camellia japonica* disease, the betaflexiviruses characterized in this study show an unusual hyper-conservation of the coat protein at aminoacidic level.

## INTRODUCTION

*Camellia japonica* is an evergreen subcanopy tree belonging to the Theaceae family, genus *Camellia.* It is the most important ornamental species in its taxonomic group (Vela et al., 2013) not only for the aesthetic beauty of its flowers but also for its role in medical and cosmetic fields: in fact, recent investigations showed its value in terms of bioactive compound content and antioxidant profile (Kim et al., 2019; Lu et al., 2019; Páscoa et al., 2019).

In Japan and in South Korea, *C. japonica* is a naturally widespread plant species, predominant in old-grown forests and islands and typically blooming from the end of January to March (Chung et al., 2003). The *C. japonica* species was first noticed in China at the end of the 17th century and imported in England, from where it diffused to Italy, which became in a very short period the main center of seed production. At the end of the 18th century this species spread and became popular also in the Americas (Hume, 1955).

Since *C. japonica*’s shrubs produce a low number of fruits that contain very few seeds (San José et al., 2016), the propagation methods in commercial nurseries rely not only on seeds, but also on hardwood cutting (preferred by Europeans and Americans) and grafting (International Camellia Society, 2019). Cutting and grafting techniques can induce the diffusion and the persistence of different kind of pathogens through plant generations but also viral transmission through seed can be possible in *C. japonica* (Liu et al., 2019).

In this regard, viral symptoms affecting camellia plants are described in literature since the late 1940: color-breaking of flowers, yellow mottle, necrotic rings and ringspots on leaves. These symptoms are recognized as typical of the Camellia leaf yellow mottle (CLYM) disease, transmissible by graft but not by sap inoculation. This disease was associated to the presence of rod-shaped viral particles (140-150 x 25-30 nm) in the cytoplasm (rarely in the nucleus) identified solely through electron microscopy (Gailhofer et al., 1988; Hiruki, 1984; Miličić, 1989). This virus was named Camellia yellow mottle virus (CYMoV) and, due to its helicoidal morphology, was initially proposed as a member of the genus *Varicosavirus* but never classified by the International Committee on Taxonomy of Viruses; its vector is still unknown (Valverde et al., 2012). In India, in 1970, another virus infecting *C. japonica* plants was discovered: it was called Tearose yellow mosaic virus (TRYMV) and was successfully transmitted to healthy plant with the aphid *Toxoptera aurantii* (Ahlawat and Sardar, 1973).

Recently, thanks to modern viral investigation techniques, such as the Next Generation Sequencing (NGS) approach, new viral species probably involved in some *C. japonica* diseases were described. In 2018 in fact, Zhang and colleagues (Zhang et al., 2018) using this method, identified a novel geminivirus called Camellia chlorotic dwarf-associated virus (CaCDaV) associated with chlorotic dwarf disease in which the affected plants display young leaves with chlorosis, deformations and V-shaped margins. A recent work allowed the association of foliar chlorotic ringspot symptom (that occurred with or without other symptoms like mottle and/or leaf variegation) with three novel viruses of the family *Betaflexiviridae*, which were detected also in seeds of diseased plants (Liu et al., 2019). Here we report a two-year investigation on the virome of Italian camellia plants showing virus-like symptoms. As initial attempts of mechanical transmission of a possible infectious agent failed, NGS analyses were performed. A complex virome was revealed, composed of a number of virus sequences belonging to the families *Fimoviridae* and *Betaflexiviridae*. Sequences were characterized and associated to two new species of the *Emaravirus* genus and 5 betaflexivirus sequences clustering with those recently characterized from samples from USA (Liu et al., 2019).

## MATERIALS AND METHODS

### Plant material and sap transmission

Plants of *Camellia japonica* showing variegation symptoms (principally on leaves, sometimes on colored flowers) were selected for sample collection from different nurseries in the area of Lake Maggiore (Piedmont, Italy) from year 2017 to 2019.

In order to understand if a putative viral etiological agent could be mechanically transmitted, leaf extracts from symptomatic plants were mechanically inoculated to a number of herbaceous test plants as already described (Roggero et al., 2002).

### Transmission electron microscopy

For negative staining, portions of infected leaves were crushed and homogenized in 0.1 M phosphate buffer, pH 7.0, containing 2% PVP. A drop of the crude extract was allowed to adsorb for 3 min on carbon and formvar-coated grids and then rinsed several times with water. Grids were negatively stained with aqueous 0.5% uranyl acetate and excess fluid was removed with filter paper.

For sections, squared pieces of about 5 mm each dimension were excised from symptomatic leaves and embedded in Epon epoxy resin (Sigma). Briefly, they were immediately sub-merged in the fixation solution (2.5% glutaraldehyde in 100 mM phosphate buffer pH 6.8), vacuum treated and then incubated over night at 4°C. Samples were rinsed three times for 5 min in 100 mM phosphate buffer pH 6.8, cut in small strips of no more than 1 mm of width and then treated as described in (Rossi et al., 2018). Ultrathin sections (70 nm in thickness) were cut using an ultra-microtome (Reichert-Jung Ultracut E, Leica Microsystems, Wetzlar, Germany), collected on formvar coated copper/palladium grids and stained for 1 min with lead citrate (Reynolds, 1963).

Observation and photographs were made with a PHILIPS CM10 TEM (Eindhoven, The Netherlands), operating at 60 kV. Micrograph films were developed, digitally acquired at high resolution with a D800 Nikon camera; images were trimmed and adjusted for brightness and contrast using GIMP 2 software.

### Isolation of viral RNA

About 1 g of symptomatic and asymptomatic camellia leaves were collected for RNA extraction, and RNA extraction was performed using the protocol described in (McGavin et al., 2012) with slight modifications. The sampled leaf tissues were extracted using 6 ml of HB buffer [0.05 M Tris/HCl (pH 8.0), 0.02 M EDTA, 0.25 M sodium sulphite, 1% polyvinylpyrrolidone, 0.02 M sodium diethyldithiocarbamate]. The homogenate of the sample was prepared using a mechanical press, by grinding the leaf tissues in specific filter bags (BIOREBA). Filtered extracts were collected and added with PEG (10%), 0.2 M NaCl and 5% Triton X-100 and then stirred for 1 h in the cold room (4°C). The resulting mixture was centrifuged for 40 min at 10.000 rpm. in a Sorvall rotor GSA; the pellet obtained after the centrifugation was resuspended in 300 microliters of 1% TE buffer [1 M Tris/HCl (pH 8.0), 0,5 M EDTA (pH 8.0)] and centrifuged again in a microfuge for 10 min at 10.650 g. The supernatant was collected avoiding to disturb the pellet and was mixed with 750 microliters of Binding buffer of Total Spectrum RNA kit (Sigma–Aldrich, Saint Louis, MO, USA). Subsequently the extraction proceeded following manufacturer instructions.

### RNAseq

The RNA samples were quantified with a NanoDrop 2000 Spectrophotometer (Thermoscientific, Waltham, MA, USA). For the first NGS analysis performed in 2018, the RNA samples extracted from symptomatic plants (listed in Table 2.) were pooled together by mixing 1 μg of RNA from each sample in a single pool. For the second NGS analyses (2019) RNAs extracted from four plants (three symptomatic and one asymptomatic) were maintained separated. RNA was sent to sequencing facilities (Macrogen, Seoul, Rep. of Korea): ribosomal RNAs (rRNA) were depleted (Ribo-ZeroTM Gold Kit,Epicentre, Madison, USA), cDNA libraries were produced (TrueSeq totalRNA sample kit, Illumina) and sequencing were carried out by an Illumina HiSeq4000 system generating paired-end sequences.

### Transcriptome assembly

The pipeline for transcriptome assembly includes 4 steps: cleaning, assembly, blasting and mapping. Reads were cleaned using BBtools (Bushnell et al., 2017), by removing adapters, artifacts, short reads and ribosomial sequences. Trinity software (version 2.3.2) (Haas et al., 2013) was used for de novo assembly of the cleaned reads. A custom viral database was used to search virus sequences in the assembled contigs via NCBI blast toolkit (version 2.8). After manual validation, the positive hits corresponding to viral sequences, were blasted against NCBInr (release October 2018) using DIAMOND (Buchfink et al., 2015). In order to obtain the number of reads mapping on each viral sequence, the viral hits were mapped on the viral contigs using bwa (Li and Durbin, 2009) and transformed with samtools (Li et al., 2009). Tablet software (Milne et al., 2016) has been used to visualize the reads mapping on viral genomic segments. For prediction of protein Open Reading Frames, ORFfinder was used with default parameters (Rombel et al., 2002).

### Confirmation of the presence in the RNA extracts of the viral contigs assembled *in-silico*

The RNA samples were retro-transcribed to cDNA at 42°C for 1h using random hexamers of the RevertAid RT Reverse Transcription Kit (Thermo Scientific, Waltham, MA, USA) following manufacturer instructions. Quantitative RT-PCR were performed using a CFX Connect™ Real-Time PCR Detection System (Bio-Rad Laboratories, Hercules, CA, USA) and iTaq™ Universal SYBR^®^ Green Supermix as previously described (Picarelli et al., 2019)

Conventional RT-PCR was performed using Phusion® High-Fidelity DNA Polymerase kit (New England Biolabs) following the manufacturer’s instructions. The PCR conditions were as follows: 30s initial denaturation at 98°C followed by 35 cycles of 10 s denaturation at 98°C, annealing 30 s at 54°C, elongation 40 s at 72°C and a final extension 5 min at 72°C.

All the primers used in these experiments are listed in supplementary Table 1.

### Phylogenetic analysis and identity/similarity matrices construction

Protein sequences coded by the putative viral fragments were used for the research of similar sequences in GenBank, and used to derive phylogenetic trees. Viral proteins were correctly aligned using MUSCLE and the alignments were processed applying the following tools: ModelFinder (Kalyaanamoorthy et al., 2017), IQ-TREE for trees reconstruction (Nguyen et al., 2014) and finally the ultrafast bootstrap (1000 replicates) (Diep Thi et al., 2018).

All the accession numbers of the proteins included in the trees are listed in Supplementary Table 2. To construct the identity/similarity matrices, the MUSCLE alignments (of every emaravirus and betaflexivirus protein) were elaborated using the online tool SIAS (Sequence Identity and Similarity) (SPAIN RESEARCH AGENCY & U.C.M. Research Office) and the sequence comparison analyses were performed applying the BLOSUM62 substitution matrix.

### Bioinformatics analysis for the identification of further fragments

In order to identify additional fragments belonging to emaraviruses the following bioinformatics strategy has been applied: i) a virus free library, derived from “healthy plant-2019”, has been used as reference to subtract the common contigs from virus infected plant libraries derived from CAM-NGS2018, CAM1-NGS2019, CAM2-NGS2019 and CAM3-NGS2019samples. ii) The remaining contigs, only present in infected libraries, have been compared, using NCBI blast toolkit, to identify contigs with high identity between the two libraries (NGS 2018-NGS 2019). iii) The list of common contigs has been blasted against NCBI nr database (version: October 2018), to remove the already known sequences. The resulting 5 candidate fragments have been mapped with bwa and confirmed by qRT-PCR as described above.

## RESULTS

### Virus-like particles associated to symptomatic *Camellia japonica* plants

Many *Camellia japonica* plants showing viral symptoms pictured in Figure 1 were reported in different sites of Piedmont (Italy): leaves displayed chlorotic ringspots (Fig.1, A), deformations (Fig.1, B), necrotic rings (Fig.1, C) and yellowing associated to necrotic rings (Fig.1, D); not only mature leaves manifested the investigated symptoms but also young-fresh leaves were affected (Fig.1, B). During the study of the symptoms, we noticed also deformations and color breaking of petals (Supplementary Figure 1.) already described in literature (Gailhofer et al., 1988; Hiruki, 1984). Negative staining of symptomatic leaves showed coiled virus-like particles as those described in Prunus by (James et al., 1999). Particles ranged from completely coiled, partially uncoiled and totally uncoiled structures (Fig. 2a, b, c, d). Completely coiled particles showed 12 loops, length of about 130 nm, an average width of 31 nm and a short extension at one or both ends (Fig. 2a). Partially uncoiled particles showed less than twelve loops and longer filamentous extensions at one or both ends of about 11 nm in diameter (Fig. 2b,c), which is the diameter observed also for totally uncoiled particles (Fig. 2d). In ultrathin sections of symptomatic leaves, spherical double-enveloped bodies, approximately 60-70 nm in diameter, were observed (Fig. 2 e, f, g).

**Figure 1.**
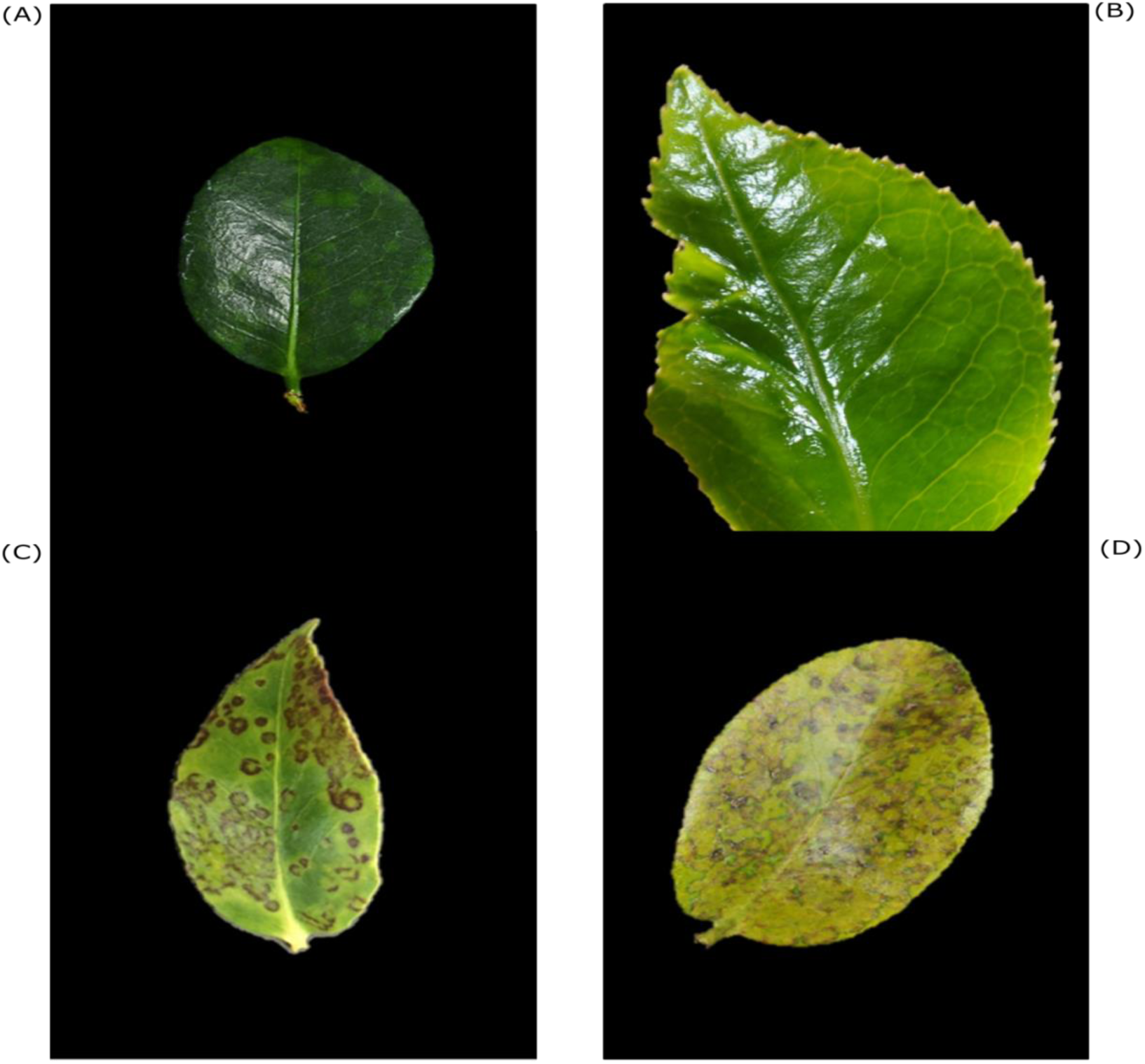
Symptoms on *Camellia japonica* leaves: chlorotic ringspots (A), malformations (B), necrotic rings (C) and yellowing with necrotic rings (D)

**Figure 2.**
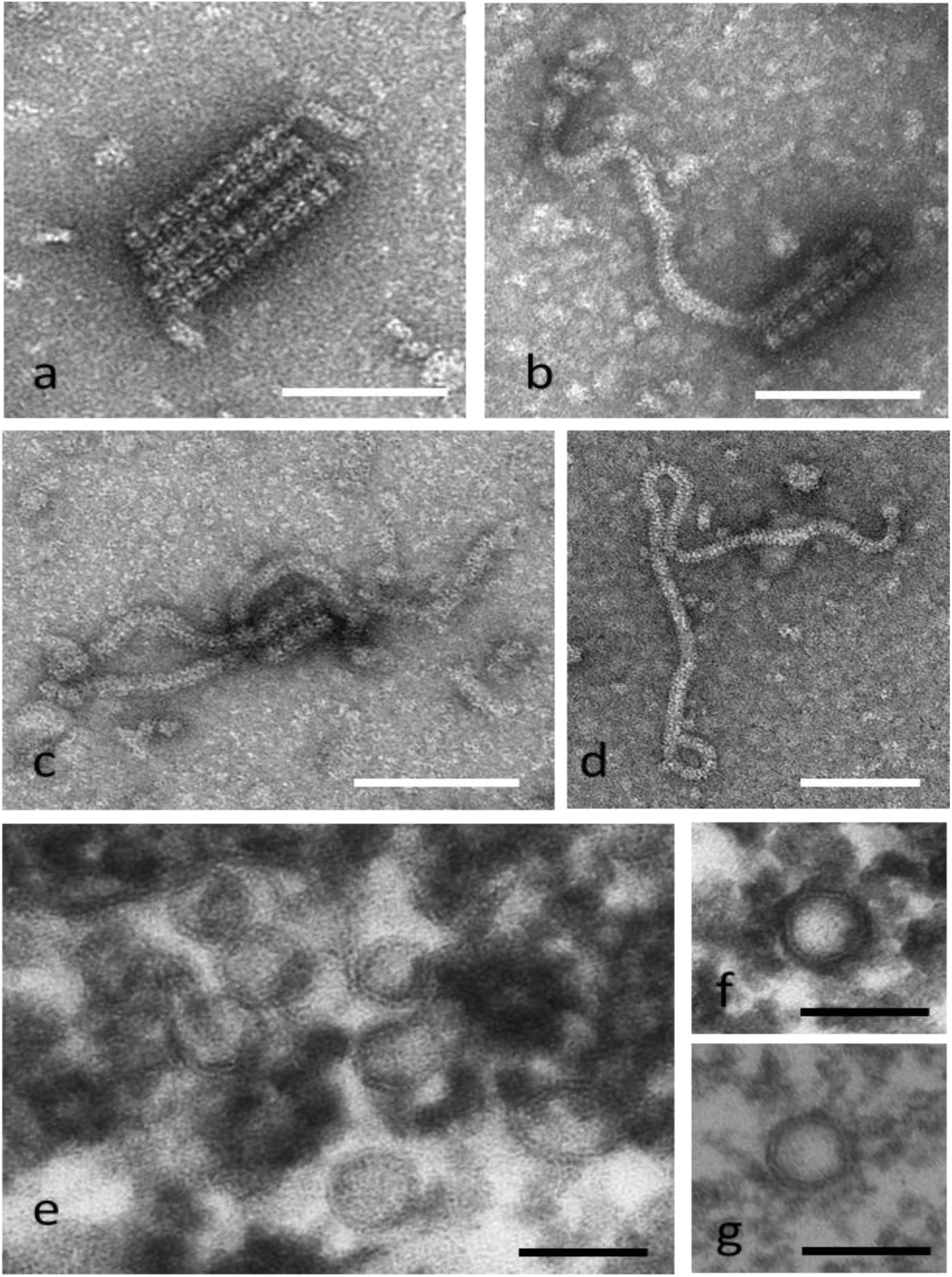
Coiled virus-like particles observed in negative staining of crude extract (a, b, c, d) and spherical double-enveloped bodies in ultrathin sections (e, f, g) of symptomatic camellia leaves. Scale bar: 100 nm

After confirming virus-like particles in the samples we tried to transmit the viruses to herbaceous healthy plants by mechanical inoculation, but without success (systemic leaves of inoculated plants were tested by specific qRT-PCR –data not shown-).

### New emaraviruses associated to symptomatic italian *Camellia japonica* plants

Two NGS analyses were performed on symptomatic *C. japonica* plants. A first one in 2018 on a pool of leaves collected at the end of 2017 (CAM-NGS2018 sample) and a second one in 2019 on four distinct samples, three from symptomatic plants (CAM1-NGS2019, CAM2-NGS2019 and CAM3-NGS2019) and one from a symptomless plant called “healthy plant 2019”. The contigs of the different viral RNAs were assembled and viral sequences belonging to different viral species were identified in all the samples (see Table 1), except for healthy plant 2019.

**Table 1.** Viruses identified in this work with the accession numbers and the mapping of the reads for every genomic segment.

| Virus name | Genome segment | Accession number | Mapping reads sample: CAM-NGS2018 | Mapping reads sample: CAM1-NGS2019 | Mapping reads sample: CAM2-NGS2019 | Mapping reads sample: CAM3-NGS2019 | Mapping reads sample: healthy plant 2019 | Contig lenght |
| --- | --- | --- | --- | --- | --- | --- | --- | --- |
| CjBV1-2018 | Genomic RNA | MN385581 | 20692 | 709 | 21166 | 37962 | 0 | 7676 |
| CjBV1-2-2019 | Genomic RNA | MN532567 | 9858 | 50 | 54694 | 110335 | 0 | 7605 |
| CjBV1-3-2019 | Genomic RNA | MN532565 | 25143 | 44081 | 5230 | 10634 | 0 | 7744 |
| CjBV2-3-2019 | Genomic RNA | MN385582 | 109385 | 68209 | 102591 | 101899 | 0 | 7173 |
| CjBV3-3-2019 | Genomic RNA | MN532566 | 197035 | 124860 | 197968 | 177282 | 0 | 7791 |
| CjEV1 | RNA1 (RdRp) | MN385573 | 31107 | 5507 | 35038 | 147274 | 0 | 7109 |
| CjEV2 | RNA1 (RdRp) | MN385577 | 17656 | 0 | 195 | 468 | 0 | 7120 |
| CjEV1 | RNA2 (GP) | MN385574 | 3530 | 1069 | 7821 | 42858 | 0 | 2054 |
| CjEV2 | RNA2 (GP) | MN385578 | 1659 | 0 | 58 | 92 | 0 | 2089 |
| CjEV1 | RNA3 (NC) | MN385575 | 4896 | 1256 | 5873 | 36763 | 0 | 1360 |
| CjEV2 | RNA3 (NC) | MN385579 | 2092 | 0 | 32 | 151 | 0 | 1316 |
| CjEV1 | RNA4 (MP) | MN385576 | 19113 | 1717 | 13070 | 78527 | 0 | 1349 |
| CjEV2 | RNA4 (MP) | MN385580 | 4414 | 0 | 62 | 158 | 0 | 1154 |
| CjEV1 | Putative RNA5 (unknown protein) | MN557024 | 1647 | 206 | 855 | 9855 | 0 | 1246 |
| CjEV1 | Putative RNA6 (unknown protein) | MN557025 | 747 | 228 | 1271 | 7879 | 0 | 1474 |
| CjEV1 | Putative RNA7 (unknown protein) | MN557026 | 5230 | 969 | 8222 | 40981 | 0 | 1297 |
| CjEV1 | Putative RNA8 (unknown protein) | MN557027 | 3129 | 413 | 3824 | 21346 | 0 | 1335 |
| CjEV1 | Putative RNA9 (unknown protein) | MN557028 | 349 | 146 | 731 | 3971 | 0 | 1155 |

Some of the sequences found in the sample CAM-NGS2018 (Supplementary Fig. 2, A) matched with emaraviruses genomic fragments that have negative stranded (-) ssRNA genomes. We were able to identify eight fragments corresponding to two RNA1, two RNA2, two RNA3 and two RNA4, supposedly belonging to two distinct emaraviruses.

The two RNA1 fragments were both 7119 nucleotides in length with a single ORF coding for putative RNA dependent RNA polymerases (named RdRp1 and RdRp2), with a predicted molecular weight of 274 kDa and 275 kDa respectively. However further investigations showed that these full length sequences were not actually present in the sample as assembled by trinity (see results below).

RNA2 segments were 2054 (accession number MN385574) and 2089 nucleotides (accession number MN385578) in length, respectively. They both code for a putative glycoprotein (GP) of 76 kDa and 76.5 kDa each, which share a 46.67% (99% of coverage) aa identity in a Blast alignment. Taken individually, they have the highest identity to the GP precursor of the emaravirus high plains wheat mosaic virus (YP_009237256.1) with percentage of 28.87% (coverage 78%) and 28.49% (73% coverage), respectively.

RNA3 segments were, respectively, 1360 nucleotides (accession number MN385575) and 1316 nucleotides (accession number MN385579) in length. Their ORFs code for putative nucleocapsid proteins (NP) with a predicted molecular weight of 33.9 kDa and 34.7 kDa respectively, which have an aa identity of 44.74% (99% of coverage) between them. MN385575 is more similar to wheat mosaic virus NP protein (AML03167.1) with an aa identity of 24.71% (coverage of 56%), whereas MN385579 has an aa identity of 24.86% (58% of coverage) with the NP protein of redbud yellow ringspot-associated emaravirus (AEO88241.1).

Finally, the two RNA4 segments were 1349 nucleotides (accession number MN385576) and 1154 nucleotides (accession number MN385580) in length and their ORFs code for a putative movement protein (MP) of 39.5 kDa and 39.8 kDa, with a percentage of aa identity between them of 75.23% (97% of coverage). Both MN385576 and MN385580 amino acidic sequences are similar to palo verde broom virus MP (AWH90178.1) with 28.44% (85% of coverage) and 28.18% of identity (84% of coverage) respectively.

In the 2019 set of samples analyzed by NGS (Table 1) four sequences matching with emaravirus genomic fragments were found. RNA2 (MN385574), RNA3 (MN385575) and RNA4 (MN385576) segments were identical to the ones already described in CAM-NGS2018. Surprisingly, RNA1 was different from both RdRp1 and RdRp2 coding sequences found in 2018 sample. This new RNA1 fragment (accession number MN385573, see Supplementary Fig. 2, B) of 7109 nucleotides in length, encodes a putative RdRp (named RdRp3), with the predicted molecular weight of 275.2 kDa and an aa identity of 32.02% (85% of coverage) with the RdRp of ti ringspot-associated emaravirus (QAB47307.1). The alignment of the protein sequence of this new ORF with the protein sequences of RdRp1 and RdRp2 previously found (in 2018 sample), showed that from aa 1 to aa 1384 it was identical to RdRp2 and that from aa 1366 to aa 2320 it was identical to RdRp1. This result suggests that RdRp3 could be the consequence of a recombination event involving RdRp1 and RdRp2 identified in the first NGS analysis (2018).

Moreover, in 2019 NGS analyses we identified five more putative viral ssRNA (-) fragments that showed a conserved and complementary short sequence of nucleotides characteristic of the emaraviruses (Mielke-Ehret and Mühlbach, 2012) to their 3’ and 5’ ends. Putative RNA5 (MN557024), RNA6 (MN557025), RNA7 (MN557026), RNA8 (MN557027) and RNA9 (MN557028) are, respectively, 1246, 1474, 1297, 1335, 1155 nucleotides in length and their ORFs encode hypothetical proteins of 21.8 kDa, 23.9 kDa, 25 kDa, 25.7 kDa, 33.7 kDa (see Table 1. and Supplementary Figure 2., C).

Only the protein coded by the ORF of putative RNA7 shares similarity with a hypothetical protein of the emaravirus wheat mosaic virus (AML03179) for a 29.30% of aa identity (98% of coverage). The proteins coded by the ORFs of putative RNA7 and putative RNA8 have an aa identity of 52.75% (100% coverage) between them.

### Putative Emaravirus recombination was not confirmed through RT-PCR

To better understand whether the RdRp3 could effectively be the result of a recombination event, we decided to carry out a specific PCR experiment using primers flanking a “transition zone” (represented in Fig.3, A) where the three RdRp1, RdRp2 and the RdRp3 coding sequences share 29 identical nucleotides. More in detail, four reactions were prepared (for the scheme, see Fig.3, A): mix 1, to amplify a fragment of 438 nucleotides of the segment encoding the putative RdRp1; mix 2, to amplify a fragment of 435 nucleotides of the segment encoding the putative RdRp2; mix 3 to amplify a fragment of 438 nucleotides of the segment encoding for the putative RdRp3; mix 4, prepared to rule out the presence of a fourth recombinant RdRp, amplifying an hypothetical fragment of 435 nucleotides. Reactions were run on two RNA samples extracted from two different plants (called Sample A, corresponding to CAM3-NGS2019, and Sample B-Silver waves see Table 2). As shown in Fig. 3 (B), bands of the expected size were obtained in both the samples with mix 3, and only in sample B with mix 4. Unexpectedly, no bands were obtained with mix 1 and 2. These results demonstrated that: i) RdRp1 and RdRp2 were the consequence of an incorrect *in-silico* assembly of the sequences obtained from the NGS analyses and were not real; ii) RdRp3 is not a recombinant version of RdRp1 and RdRp2. Moreover, mix 4 highlighted the presence of a new RdRp sequence (named RdRp4) in camellia samples showing a first part of the fragment (from aa 1 to aa 1384) identical to RdRp1 sequence and a second part (from aa 1366 to aa 2324) identical to RdRp2 sequence. The new RNA1 coding for RdRp4 (accession number MN385577; Fig.4) is 7120 nucleotides in length and its ORF encodes a protein of 274.5 kDa. RdRp4 shares an aa identity of 59.64% (coverage of 99%) with RdRp3 and it is similar to a RNA replicase p1 of Pistacia emaravirus (QAR18002.1) (aa identity of 33.09%; coverage of 85%).

**Table 2.** Camellia plants (*Camellia japonica, Camellia higo* and *Camellia hybrid*) analyzed in this study and the diagnoses based on qRT-PCR.

| Cultivar | Source | Virus |
| --- | --- | --- |
| *Camellia japonica*, Drama girl | Villa Giuseppina<br>Verbania, Piedmont | CjBV2 |
| *Camellia higo*, Hiodoshi | Villa Giuseppina<br>Verbania, Piedmont | none |
| *Camellia higo*, Shiro osaraku | Villa Giuseppina<br>Verbania, Piedmont | CjBV2 |
| *Camellia japonica*, Margaret Davis | Villa Giuseppina<br>Verbania, Piedmont | CjBV1, CjBV2 |
| *Camellia japonica*, Prof Giovanni Santarelli | Villa Giuseppina<br>Verbania, Piedmont | none |
| *Camellia hybrid*, Mary Phoebe taylor | Villa Giuseppina<br>Verbania, Piedmont | CjBV2 |
| *Camellia japonica*, Shiro Kinjo | Villa Giuseppina<br>Verbania, Piedmont | none |
| *Camellia japonica*, Chubu tsukimi guruma | Villa Giuseppina<br>Verbania, Piedmont | CjBV1,CjBV2 |
| *Camellia hybrid*, Valley Knudsen | Villa Giuseppina<br>Verbania, Piedmont | none |
| *Camellia hybrid*, Dr Clifford Parks | Villa Giuseppina<br>Verbania, Piedmont | CjBV2 |
| *Camellia japonica*, Kirin-no-homare | Villa Giuseppina<br>Verbania, Piedmont | none |
| *Camellia japonica X* | Villa Giuseppina<br>Verbania, Piedmont | CjBV2 |
| *Camellia japonica*, Nobilissima | Nursery 1, Verbania Piedmont | CjBV2 |
| *Camellia japonica*, Nuccio's jewel 6 | Nursery 1, Verbania Piedmont | CjEV1, CjEV2, CjBV1, CjBV2 |
| *Camellia japonica*, Nuccio's jewel 10 | Nursery 1, Verbania Piedmont | CjEV1, CjEV2, CjBV1, CjBV2 |
| *Camellia japonica*, Nuccio's jewel 22 | Nursery 1, Verbania Piedmont | CjEV1, CjEV2, CjBV1, CjBV2 |
| *Camellia japonica*, Silver waves<br>(Sample B) | Nursery 1, Verbania Piedmont | CjEV1, CjEV2, CjBV1, CjBV2 |
| *Camellia japonica*, California * 1<br>(sample CAM-NGS2018) | Nursery 2, Verbania Piedmont | CjEV1, CjEV2, CjBV1, CjBV2 |
| *Camellia japonica*, California * 2<br>(sample CAM-NGS2018) | Nursery 2, Verbania Piedmont | CjEV1, CjBV1 |
| *Camellia japonica*, California *3<br>(sample CAM-NGS2018) | Nursery 2, Verbania Piedmont | CjEV1, CjEV2, CjBV1 |
| *Camellia japonica*, California 3 | Nursery 2, Verbania Piedmont | CjEV1, CjEV2, CjBV1 |
| *Camellia japonica*, California 4 | Nursery 2, Verbania Piedmont | CjEV1, CjBV1, CjBV2 |
| *Camellia japonica*, California 1 | Nursery 2, Verbania Piedmont | none |
| *Camellia japonica*, California 2 | Nursery 2, Verbania Piedmont | none |
| *Camellia japonica 1* | Turin, Piedmont | CjBV1,CjBV2 |
| *Camellia japonica 2* | Turin, Piedmont | CjBV2 |
| *Camellia japonica 3* | Turin, Piedmont | CjBV2 |
| *Camellia japonica 4* | Turin, Piedmont | CjBV2 |
| *Camellia japonica*, California *<br>(sample CAM1-NGS2019) 1 | Nursery 2, Verbania Piedmont | CjEV1, CjBV1,CjBV2 |
| *Camellia japonica*, California *<br>(sample CAM2-NGS2019) 2 | Nursery 2, Verbania Piedmont | CjEV1, CjBV1, CjBV2 |
| *Camellia japonica*, California *<br>(sample CAM3-NGS2019) (Sample A) 3 | Nursery 2, Verbania Piedmont | CjEV1, CjBV1, CjBV2 |
| *Camellia japonica*, General Coletti 4 | Flower shop,Turin, Piedmont | CjBV1, CjBV2 |
| *Camellia japonica*, RL Wheeler*<br>( sample healthy plant 2019) | Flower shop,Turin, Piedmont | none |
| *Camellia japonica*, Dr Burnside 7 | Flower shop,Turin, Piedmont | CjBV2 |
| *Camellia japonica*, Margaret Wells 6 | Flower shop,Turin, Piedmont | CjBV1, CjBV2 |

\* Plants analyzed by NGS; in bolt: symptomatic plants (one or more putative viral symptoms). When the variety is not written, is unknown.

**Figure 3.**
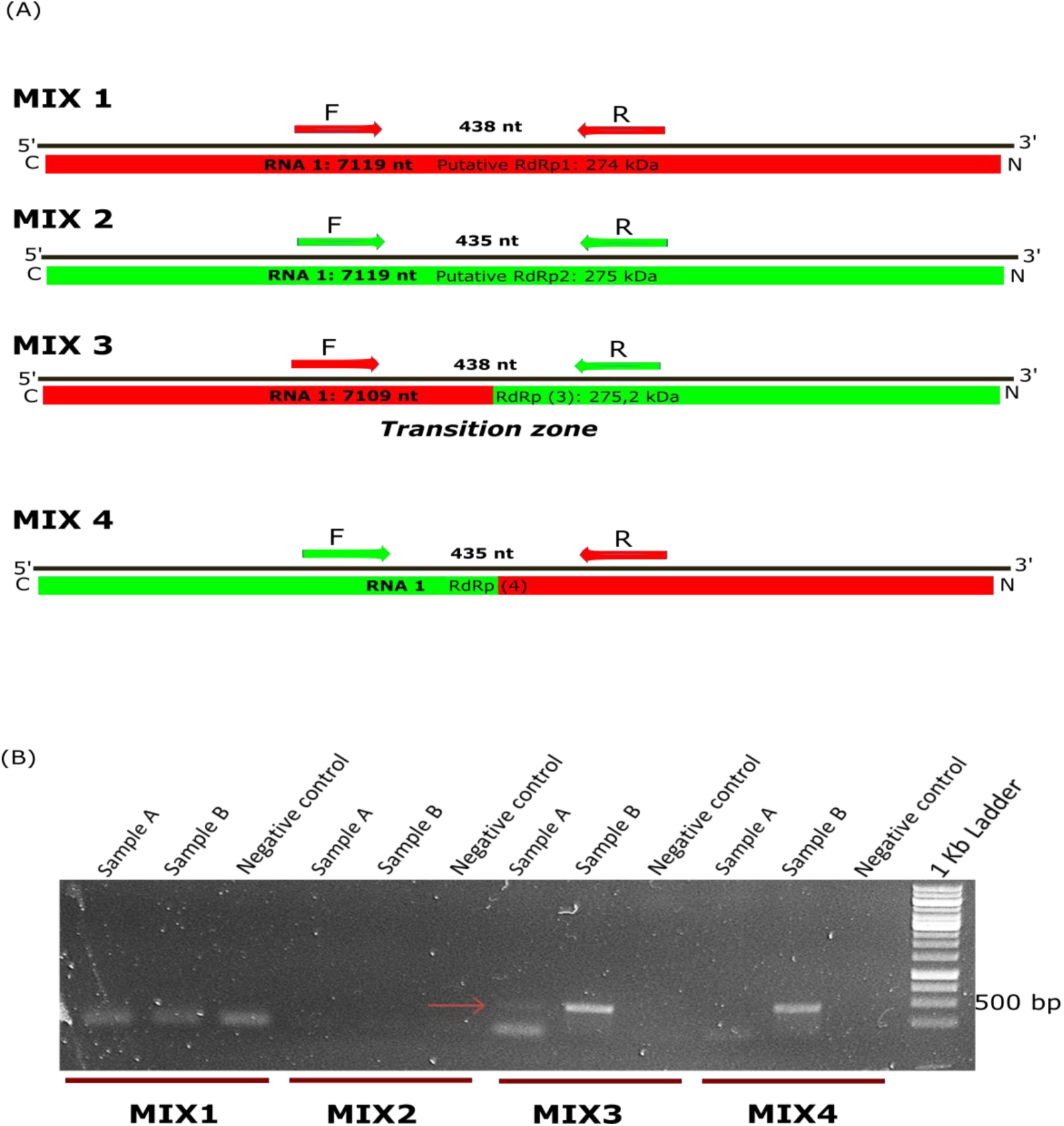
Schematic overview of the detection of the emaravirus RdRp transition zone (A), colored rectangles are the open reading frame (ORF) while black lines represent the genome segments of the RdRp: MIX1 contains primer specifically designed across the transition zone on the *in-silico* assembled RdRp1 encoding sequence, MIX2 includes primers specifically designed across the transition zone on RdRp2 encoding sequence, MIX3 contains a primer F specifically designed on RdRp1 sequence and a primer R specifically designed on RdRp2 sequence, MIX4 contains a primer F specifically designed on RdRp2 sequence and a primer R specifically designed on RdRp1 sequence; agarose gel (B) showing the amplification of the RdRp transition zone in two different samples (Sample A and Sample B) using the four MIX represented in (A), the red arrow indicates the positive band obtained in Sample A for the MIX3 (designed for the detection of RdRp 3, accession number: MN385573). Negative control= water, RdRp= RNA dependent RNA polymerase, nt= nucleotides, F= Forward primer, R= Reverse primer

**Figure 4.**
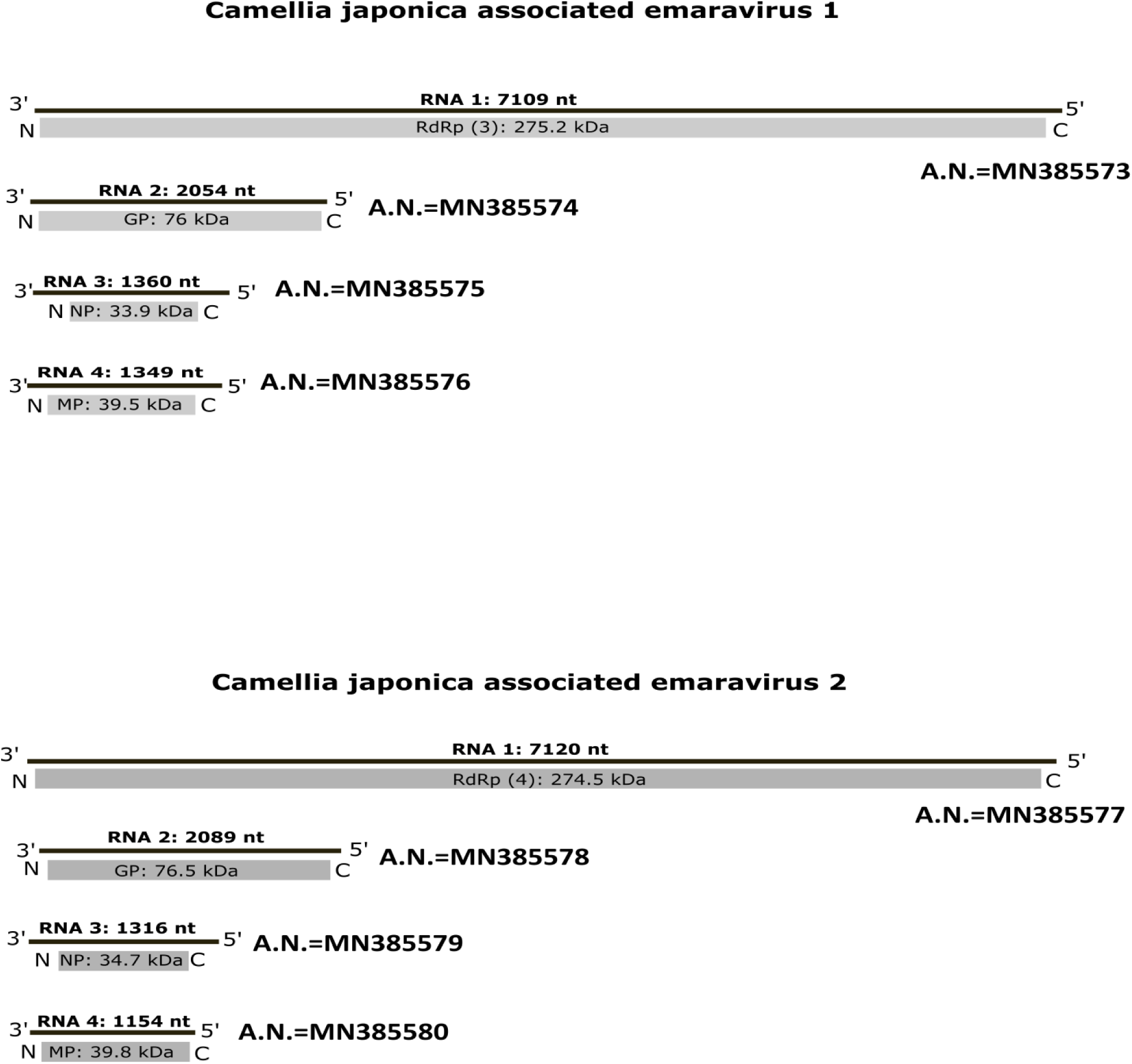
Camellia japonica associated emaravirus 1 and 2 (CjEV1 and 2) segmented genomes. A.N.= GenBank accession number

### Distribution of the genomic fragments in the CjEVs genomes

Once clarified the identification of all the eight principal emaravirus genomic fragments, the goal was to correctly associate every genomic fragment to the genomes of each of the two Camellia japonica associated emaraviruses (named CjEV1 and CjEV2).

The mappings of the reads for all the samples (see Table 1.), clearly showed that the three samples analyzed in 2019 (CAM1-NGS2019, CAM2-NGS2019, CAM3-NGS2019) were infected only by one of the two CjEV, composed of the sequences RNA1 (MN385573), RNA2 (MN385574), RNA3 (MN385575), RNA4 (MN385576), that we called CjEV1 (Fig. 4). Quantitative RT-PCR analyses using primers designed on every emaravirus RNA fragments confirmed that Sample A (corresponding to CAM3-NGS2019) was infected only by CjEV1. Sample B instead (as well as sample CAM-NGS2018) was infected by both CjEV1 and CjEV2. Therefore CjEV2 genome is formed by RNA1 (MN385577), RNA2 (MN385578), RNA3 (MN385579) and RNA4 (MN385580) (Fig. 4). Indeed none or only few reads mapped for the RNA1 (MN385577), RNA2 (MN385578), RNA3 (MN385579) and RNA4 (MN385580) in 2019 samples that only were infected with CjEV1. Further qRT-PCR analyses permitted to associate the extra putative emaraviruses RNA5 (MN557024), RNA6 (MN557025), RNA7 (MN557026), RNA8 (MN557027) and RNA9 (MN557028) to the CjEV1 genome.

### Five betaflexivirus isolates associated to symptomatic Italian *Camellia japonica* plants

NGS analyses showed also the presence of betaflexiviruses-related sequences in all the symptomatic camellia samples from both 2018 and 2019. In particular, five single strand positive RNA viral genomic sequences were identified and were associated to five viral isolates tentatively named Camellia japonica associated betaflexi-virus 1 isolate 2018 (CjBV1-2018), -virus 1 isolate CAM2-NGS2019 (CjBV1-2-2019), -virus 1 isolate CAM3-NGS2019 (CjBV1-3-2019), -virus 2 isolate CAM3-NGS2019 (CjBV2-3-2019), -virus 3 isolate CAM3-NGS2019 (CjBV2-3-2019). The mapping of reads on each isolate in each sample are shown in Table 1. Isolate CjBV1-2018 has a genomic sequence of 7676 nucleotides in length (accession number MN385581, see Fig. 5, A), in which three ORFs could be identified, coding for a putative RdRp of 226 kDa, a putative MP of 48.2 kDa and a putative CP protein of 25.1 kDa, respectively. These putative protein sequences, analyzed by a Blast search, showed similarity to the RdRp, MP and CP of Camellia ringspot associated virus 1 (Liu et al., 2019) with an identity score for each protein of 81.74%, (QEJ80622) (coverage 100%), 93.64% (QEJ80623) (coverage 100%) and 100% (QEJ80624), respectively.

**Figure 5.**
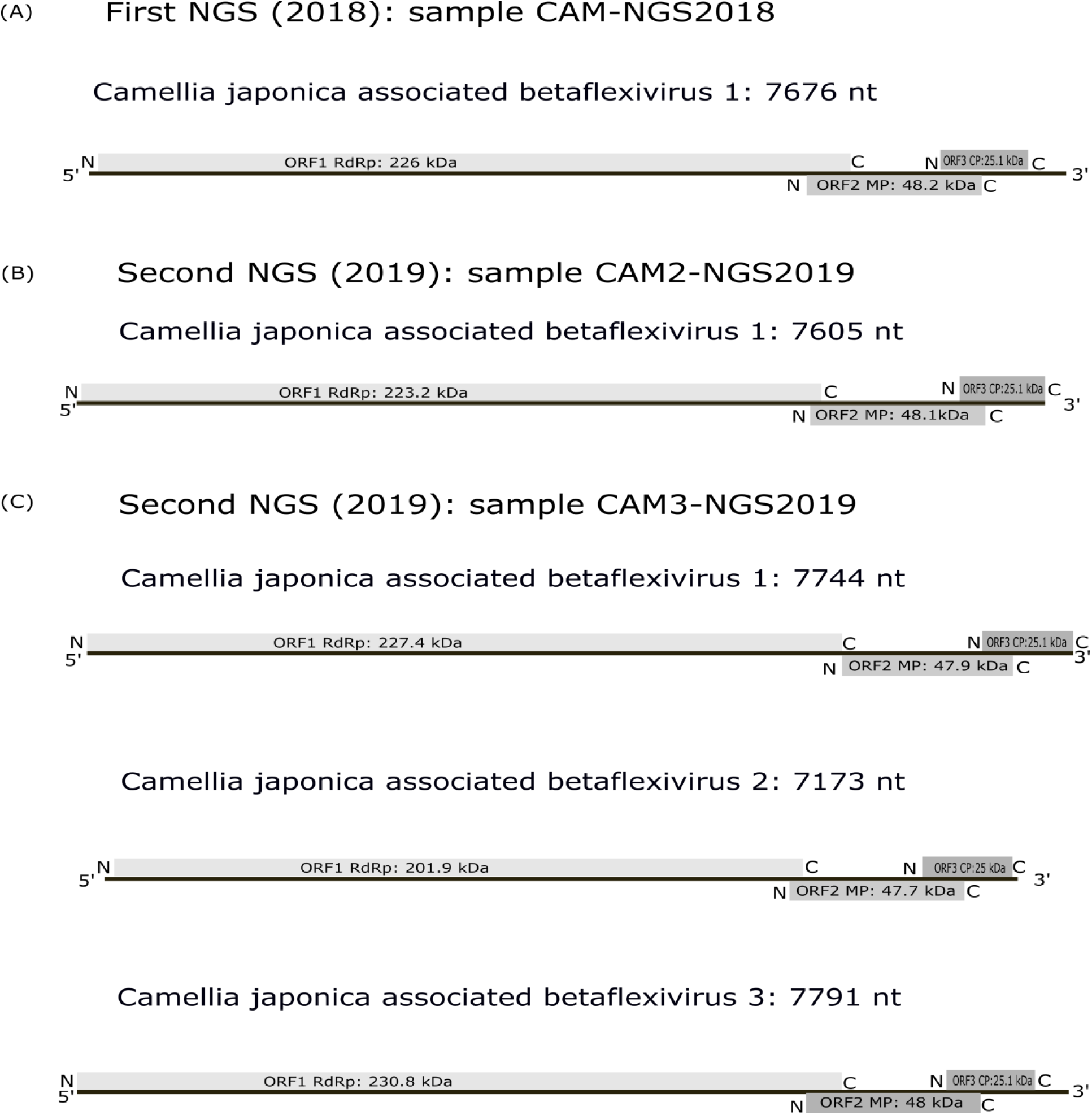
Schematic representation of betaflexivirus genomes found in the two NGS analyses. Sample CAM-NGS2018: Camellia japonica associated betaflexivirus 1 genome representation (A), Camellia japonica associated betaflexivirus 1 genome identified in the sample CAM2-NGS2019 (B), genome representations of Camellia japonica associated betaflexivirus 1, 2 and 3 found in the sample CAM3-NGS2019 (C). nt=nucleotides.

CjBV1-2-2019 has a genome of 7605 nucleotides in length (accession number MN532567) (Fig. 5, B). This sequence contains three main ORFs: incomplete ORF1, that codes for a putative RdRp of 223.2 kDa, ORF2 that encodes a putative MP of 48.1 kDa and ORF3 that encodes a putative CP protein of 25.1 kDa. Also these putative proteins are similar to the Camellia ringspot associated virus 1 (Liu et al., 2019), with an identity percentage of 81.60% (coverage 100%) for RdRp aa sequence (QEJ80622), 91.59% (100% of coverage) for MP (QEJ80623), and 99.10% (100% of coverage) for CP (QEJ80624).

CjBV1-3-2019 isolate (accession number MN532565) has a genome of 7744 nucleotides in length organized in three ORFs. ORF1 encodes a putative RdRp of 227.4 kDa, ORF2 codes for a putative MP of 47.9 kDa and ORF3 encodes a putative CP of 25.1 kDa (Fig.5, C). Again, these proteins are similar to RdRp, MP and CP of Camellia ringspot associated virus 1 proteins (Liu et al., 2019) with an identity percentage of 97.55% (100% of coverage) (QEJ80622), 98.41% (coverage of 100%) (QEJ80623), and 99.10% (100% of coverage) (QEJ80624) respectively.

CjBV2-3-2019 (accession number MN385582, Fig.5, C) genome is 7173 nucleotides long and it is formed by three main ORFs. ORF1 codes for a putative RdRp of 201.9 kDa, ORF2 encodes a putative MP of 47.7 kDa, and ORF3 codes a putative CP of 25 kDa, which resulted similar to the replicase (identity of 90.77%, coverage 100%), the MP (91.14% of identity, coverage 100%) and the CP (99.55% of identity, 100% coverage) of Camellia ringspot associated virus 2 (accession numbers QEJ80628, QEJ80629 and QEJ80627, respectively) (Liu et al., 2019).

CjBV3-3-2019 isolate (accession number MN532566, Fig.5, C) has a genome 7791 nucleotides long that contains three ORFs. ORF1 encodes a putative RdRp of 230.8 kDa, ORF2 a putative MP of 48 kDa and an ORF3 a putative CP of 25.1 kDa. The three putative proteins are, as those of CjBV2-3-2019, similar to RdRp, MP and CP of Camellia ringspot associated virus 2 (91.36% identity for RdRp (QEJ80625), 90.68% identity for MP (QEJ80626), 99.10% identity for CP (QEJ80627) (all 100% coverage).

Analyzing the aa sequences of the encoded proteins of the five CjBVs and comparing the identity percentages (see Supplementary Table 3.), it is possible to notice that CjBV1-2018, CjBV1-2-2019 and CjBV1-3-2019 have aa identity values over 80% for the three proteins (RdRp, MP,CP) when compared among them and the same is true also for CjBV2-3-2019 and CjBV3-3-2019 amino acid sequences. Moreover, when these two groups of viruses (composed one by CjBV1-2018, CjBV1-2-2019, CjBV1-3-2019 and the other by CjBV2-3-2019, CjBV3-3-2019) are compared one with the other, the aa identity values are inferior to 80% for RdRp and MP, but over 80% for the CP. Comparing the RdRp, MP and CP of each CjBV with the proteins of others betaflexiviruses the values are always inferior to 80%.

### Phylogenetic analyses

In order to frame the identified viruses in taxonomic groups and to define their possible evolutionary history, the putative amino acid sequences of the two CjEVs RdRp, GP, NP and MP proteins were aligned to those of other emaravirus protein sequences to produce phylogenetic trees (Fig. 6) In all the phylogenetic analyses, CjEV1 and CjEV2 cluster together and form a separate branch. When the RdRp, GP, MP sequences where considered, the topology of the phylogenetic trees differed only slightly and in all cases CjEVs share a common ancestor with a group of other members of the genus *Emaravirus* composed of Palo verde broom virus, high plains wheat mosaic virus, wheat mosaic virus, ti ringspot-associated emaravirus, raspberry leaf blotch emaravirus, and jujube yellow mottle-associated virus. In the case of NP sequences, the tree topology was different and the CjEVs branch clusters with a different group of emarviruses (including the species representative European mountain ash ringspot-associated emaravirus), even though such association is not supported by a bootstrap value over 70%.

**Figure 6.**
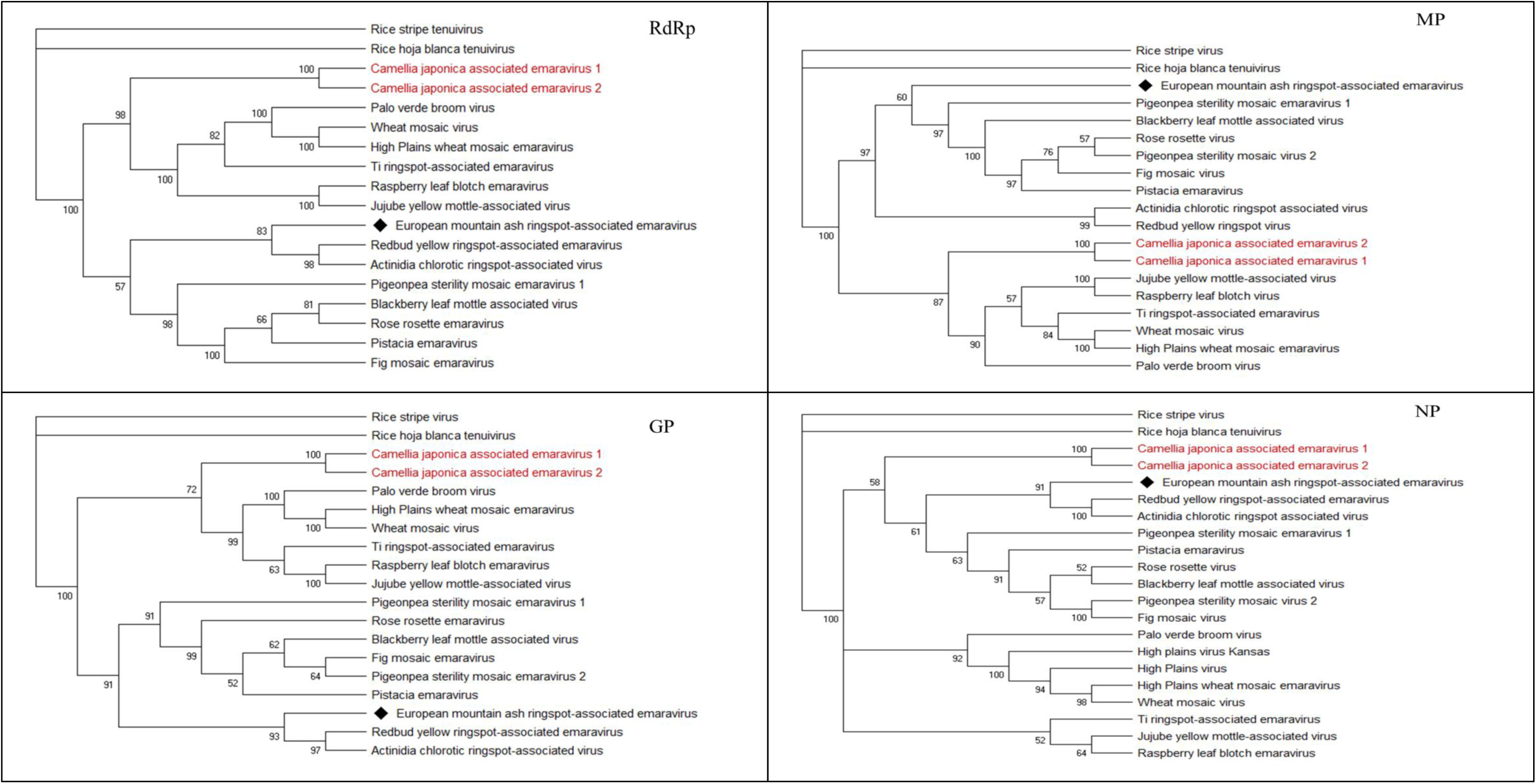
Phylogenetic placement of Camellia japonica associated emaravirus 1 and 2. Amino acids sequences of RNA-dependent RNA polymerases (RdRPs), Glycoproteins (GPs), nucleocapsid proteins (NPs) and movement proteins (MPs) were aligned with MUSCLE and then phylogenetic trees were produced using the maximum likelihood methodology in IQ-TREE software. Each branch reports numbers that represent statistical support based on bootstrap analysis (1000 replicates). The viruses identified in this work are written in red. The species representative of the emaraviruses group (ICTV taxonomy) is marked with a black diamond. The predictive models used for each phylogenetic tree are: LG+F+I+G4 (RdRp), LG+F+G4 (NP), LG+F+I+G4 (MP), WAG+F+G4 (GP)

**Figure 7.**
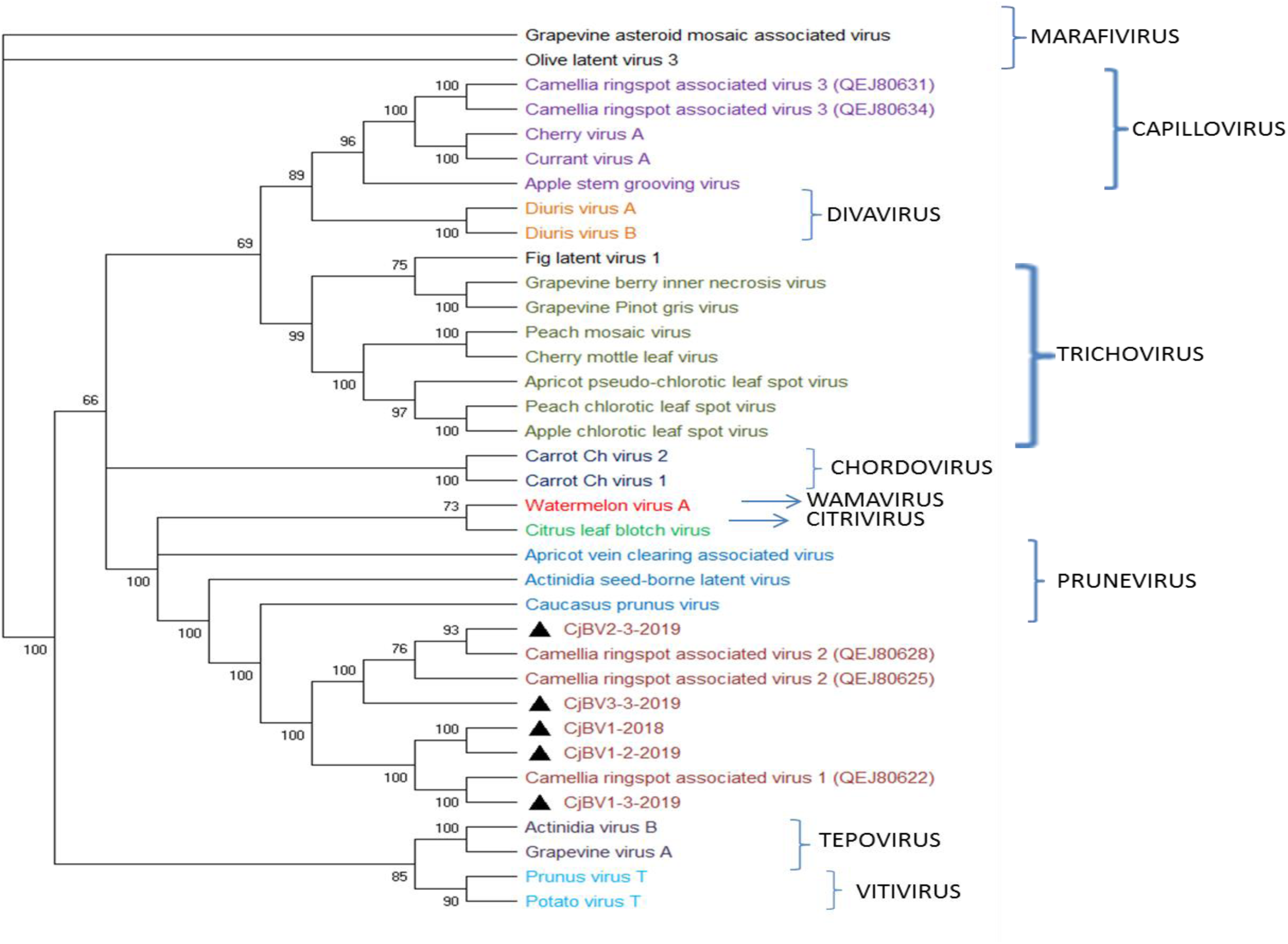
Phylogenetic placement of all Camellia japonica associated betaflexiviruses. Amino acids sequences of RNA-dependent RNA polymerases (RdRps) were aligned with MUSCLE and then phylogeny was derived using the maximum likelihood methodology in IQ-TREE software. The statistical support based on bootstrap analysis (1000 replicates) is summarized in the numbers on the branches. Viruses identified in this work are marked by black triangles. The predictive model used for the phylogenetic tree is VT+F+I+G4

The same phylogenetic analysis was also performed for the proteins coded by the three ORFs of the Camellia japonica associated betaflexiviruses (RdRp, MP and CP) (Figure 6 and Supplementary Figure 3). All the proteins of CjBV1-2018, CjBV1-2-2019 and CjBV1-3-2019 isolates cluster with the proteins of Camellia ringspot associated virus 1 (MK050792) (Liu et al., 2019) while the proteins of CjBV2-3-2019 and CjBV3-3-2019 isolates cluster with the ones of the Camellia ringspot associated virus 2 isolate CJ5-6003 (MK050794) (CRSaV-2) and Camellia ringspot associated virus 2 isolate CJ5-2013 (MK050793). Together with Camellia ringspot associated virus 1 and Camellia ringspot associated virus 2 (Liu et al., 2019), all CjBVs, form a group of viruses close, but separated from the existing members of the genus *Prunevirus*.

### Detection of the newly found viruses in Camellia samples

In order to investigate the presence of the newly identified viruses in a wider range of camellia plants, and try to associate specific virus presence with symptoms, 35 plants were analyzed through qRT-PCR. Ten plants were asymptomatic and 25 plants showed different degree of leaf variegation disease (Table 2 and Supplementary Fig.4). Reactions were performed using primers amplifying specifically the two emaraviruses species (all the single genome segments of CjEV1 and 2) and generic primers designed to amplify specifically all the sequences of each of the two betaflexivirus groups (group 1 formed by CjBV1-2018, CjBV1-2-2019, CjBV1-3-2019 and group 2 composed by CjBV2-3-2019, CjBV3-3-2019 called, respectively, CjBV1 and CjBV2 in Table 2.).

The amplifications confirmed the presence of both emaraviruses and both betaflexiviruses in 2018 sample set and the presence of both betaflexiviruses and only one emaravirus (CjEV1) in samples collected in 2019. Of the ten asymptomatic plants, four resulted negative, while six were positive for betaflexiviruses (either CjBV2 only or both). No asymptomatic plant was positive for emaraviruses. Regarding the 25 symptomatic plants, four resulted negative for all the virus we tested.

Betaflexiviruses were present in all the other 21 symptomatic plants, either CjBV2 only or both. Only the samples coming from the nurseries of Verbania Piedmont were positive also for the emaraviruses (either both or CjEV1 only).

## DISCUSSION

In this work the virome of symptomatic camellia plants sampled from 2017 (sequenced in 2018) and 2019 in Lake Maggiore in Italy was investigated. Plants showed different degrees of leaf variegation disease resembling in some cases to leaf yellow mottle and in other cases to ringspot disease. Leaf yellow mottle disease is known since 1940, however, knowledge on the pathogenic agent is still scarce and for many decades it relied only on symptoms observations and cytopathology of infected cells. Hiruki (1984) and Gailhofer et al. (1988) associated the occurrence of the disease to rod shaped particles observed in epidermal and mesophyll cells, but they were not able to mechanically transmit the causal agent. Very recently, microscopic observation and a high throughput analyses associated filamentous particles and betaflexivirus sequences to camellia foliar chlorotic and necrotic ringspots (Liu et al, 2019).

In our study, a next generation sequence approach allowed us to discover two new sequences belonging to the *Emaravirus* genus of the *Fimoviridae* family (CjEV1 and CjEV2) and five sequences belonging to the *Betaflexiviridae* family (CjBV1-2018, CjBV1-2-2019, CjBV1-3-2019, CjBV2-3-2019, CjBV2-3-2019) in Italian symptomatic camellia plants.

CjEV1 and CjEV2 are the first emaraviruses associated to camellia symptomatic plants. The genus *Emaravirus* is in the *Fimoviridae* family of the order *Bunyavirales*, whose members are plant viruses with segmented, linear, single-stranded, negative-sense RNA genomes. They are distantly related to orthotospoviruses and orthobunyaviruses (Elbeaino et al., 2018). The genus *Emaravirus* was recently established after the discovery of the *European mountain ash ringspot-associated emaravirus* (EMARaV), which is the type species, and includes also the species *Fig mosaic virus* FMV, *Rose rosette virus* RRV, *Raspberry leaf blotch virus* RLBV (Mielke-Ehret and Mühlbach, 2012). Emaraviruses have multipartite genomes organized in 4 to 8 segments of negative sense ssRNA and induce characteristic cytopathologies in their host plants, including the presence of double membrane-bound bodies (80-200 nm) in the cytoplasm of the virus-infected cells. In sections of camellia symptomatic leaves we could observe spherical double-enveloped bodies resembling those described in association with emaravirus infections (Zheng et al., 2017): however, in our case, the bodies were smaller than the expected size, since they measured approximately 60-70 nm in diameter. The *in-silico* assembly of the emaravirus sequences associated to our camellia symptomatic leaves has been particularly challenging, since we faced the need of performing a specific RT-PCR, to clarify which were the real sequences present in the samples infected by each or both emaraviruses. In fact, we found out that the RdRp1 and RdRp2 identified after the first NGS analysis were not correctly assembled, since Trinity software assembled reads that were not contiguous because of a transition region where the two RNA have a common nt sequence. Cases in which parts of viral genomes were missed or inverted are not uncommon (Hunt et al., 2015) and we demonstrated one more time that automatically assembled sequences must always be confirmed, particularly in mixed infections. Eventually we discovered the presence of two emaraviruses, CjEV1 and CjEV2, each with a core of 4 (-) ssRNA fragments, coding for RdRp (RNA1), GP (RNA2), NP (RNA3) and MP (RNA4). It is noteworthy that the four phylogenetic trees obtained for each protein, show a clear isolation of camellia RdRp, GP, NP and MP from all the other known emaraviruses proteins. This data was confirmed by the aa identity values showed in Supplementary Table 3.: according to the demarcation criteria necessary for the definition of new species - protein sequences differing more than 25% - (Elbeaino et al., 2018), CjEV1 and CjEV2 can be considered two new species. In fact, all the aa identity values resulting from the comparison of the protein sequences (RdRp, GP, NP, MP) between them and with the other emaraviruses homologous proteins were always lower than 75%. CjEV1 and CjEV2 are probably going to inhabit their own evolutionary niche in the genus *Emaravirus* which is rapidly growing. Interestingly, CjEV1 and CjEV2 NPs seem not to share the same common ancestor of the RdRp, GP and MP. This fact can be an evidence of a possible reassortment event happened in the cluster of emaravirus RNA segments maybe during a multiple infection of a *Camellia japonica* plant. Indeed, reassortment is a characteristic of viruses with segmented genomes and it is a possible way to generate new combination of segments better adapted to specific selective pressures (Margaria et al., 2015; Rastgou et al., 2009; Simon-Loriere and Holmes, 2011).

The CjBVs betaflexiviruses identified in this work share a common ancestor with the recently identified Camellia ringspot associated virus 1 (CRSaV-1), Camellia ringspot associated virus 2 (CRSaV-2) isolate CJ5-6003 (MK050794) and Camellia ringspot associated virus 2 isolate CJ5-2013 (MK050793) (Liu et al, 2019). On the base of the criteria for the definition of a new species in the family *Betaflexiviridae* of 80% aa identity in the replicase or CP genes products (Adams et al., 2012), we could identify two distinct putative viral species: CjBV1, which includes isolates CjBV1-2018, CjBV1-2-2019 and CjBV1-3-2019, associated to the previously described CRSaV-1 s, and CjBV2 that includes isolates CjBV2-3-2019 and CjBV3-3-2019, associated to the previously described CRSaV-2 group of isolates.

As already noticed by Liu and colleagues (2019), comparing the aa identity of the three proteins (RdRp, MP and CP) inside the groups of betaflexiviruses identified in Italian and American camellia plant isolates, the coat protein is always the most conserved one, (see Supplementary Table 3., for the values). Ma and colleagues (Ma et al., 2019) recently demonstrated that the CP of the Apple stem pitting virus (ASPV) (a member of the *Betaflexiviridae* family, genus *Foveavirus*) not only fulfills a protective role, encapsidating the viral genome and preserving it from the degradation but it is also involved in viral suppression of RNA silencing (VSR), one of the first lines of defense of the plant against viral attacks. This VSR property seems to be conserved among different CP variants which also have different abilities to aggregate *in vivo* in *N. benthamiana* and to cause the appearance of different symptoms in *N. occidentalis* (Ma et al., 2019). In light of this, the fact that the CP protein is so highly conserved among the group of camellia betaflexiviruses, could be linked to its role in symptoms development and in VSR in this ornamental plant, role to be explored through future studies. Our microscope observation never showed the presence of filamentous virus as the one described in Liu et al. (2019). Nevertheless, initially, the viral like particles observed in negative staining were ascribed to betaflexiviruses: in particular, the uncoiled form (Fig. 2d) resembled the Tricovirus *Apple chlorotic leaf spot virus* or the Vitivirus *Grapevine virus A* (ICTV). However, we could not find any mention in literature of betaflexiviruses forming coiled structures like the one we discovered. At the same time, we cannot exclude that those formations are related not to betaflexiviruses but to emaraviruses nucleocapsids. Further analyses are needed to elucidate the nature of those viral like particles.

During our study, we observed plants manifesting viral symptoms infected by both emaraviruses and betaflexiviridae or by betaflexiviridae only. At the same time, some symptomatic plant resulted negative for emaraviruses and betaflexivirus and plant apparently without symptoms hosted both or only one betaflexivirus but never the emaraviruses. Because camellia-infecting betaflexiviruses were found in asymptomatic plants in our work and also by Liu and colleagues (2019), it could be possible that they mostly act as cryptic viruses (Boccardo et al., 1987), but in some case they can persist also in symptomatic plants possibly hosting other yet uncharacterized viruses. This complex scenario confirm the difficulty of correlating symptoms and infectious agent. This survey should be repeated with more plants, to produce a statistical critical study of the infections, useful to better understand the dynamics in the interactions of different viral species in camellia plant and their linkage to the symptoms.

To reach this goal, first of all, an interesting possibility will be produce infective clones for CjEV1 and CjEV2 as already described by Pang and colleagues (Pang et al., 2019) that developed an innovative reverse genetic system to study the emaravirus RRV: they produced an infectious virus clone from a cDNA copy of the viral genome of RRV and in this way were able to demonstrate directly the progression of the viral disease with all its characteristic symptoms induced by RRV in Agro-infiltrated *Arabidopsis thaliana*, *Nicotiana benthamiana* and rose plants; in future this technique could be applied also to study the emaraviruses affecting *Camellia japonica* plants and to clarify if a correlation exists between the emaravirus infection and the symptoms.

These experiments could be determinant in the definition of the symptoms induced by every single virus found in the camellia virome and to understand if the disease is the result of a single or multiple viral infection.

Another important aspect of the infection cycle is the transmissibility of the emaraviruses CjEV1 and CjEV2: because many emaraviruses are transmitted by eriophyid mites (Mielke-Ehret and Mühlbach, 2012) and eriophyid mites (*Acaphylla steinwedeni, Calacarus carinatus, Cosetacus camelliae*) (Keifer, 1982) are a main threat to *Camellia japonica* plants in many environments: tests of transmissibility with eriophyid mites will be carried out to clarify if these are effectively, the vectors. To conclude, our work evidenced the existence of a complex virome in symptomatic *Camellia japonica* plants and, in particular identified two new species of *Emaravirus* genus that, at the moment, counts only 9 classified members (Elbeaino et al., 2018). Future research will be focused on clarifying the virus-symptom correlations and virus transmissibility in order to contain and eradicate Camellia plant diseases that endanger the survival and the varieties conservation of this economically important plant.

## ACKNOWLEDGMENTS

The authors would like to thank the precious technical support of Caterina Perrone and Riccardo Lenzi for their help with mechanical inoculation experiment and with the set up of the viral particles purification protocol. Moreover the authors are grateful to Gianni Morandi and Paolo Zacchera (Compagnia del Lago, Villa Giuseppina) for their availability and kindness in providing the samples and plants needed for the research.

